# ChIP-BIT2: a software tool to detect weak binding events using a Bayesian integration approach

**DOI:** 10.1101/260869

**Authors:** Xi Chen, Xu Shi, Leena Hilakivi-Clarke, Robert Clarke, Tian-Li Wang, Jianhua Xuan

## Abstract

Transcription factor binding events play important functional roles in gene regulation. It is, however, a challenging task to detect weak binding events since the ambiguity in differentiation of weak binding signals from background signals. We present a software package, ChIP-BIT2, to identify weak binding events using a Bayesian integration approach. By integrating signals from sample and input ChIP-seq data, ChIP-BIT2 can detect both strong and weak binding events at gene promoter, enhancer or the whole genome effectively. The ChIP-BIT2 package has been extensively tested on ChIP-seq data, demonstrating its wide applicability in ChIP-seq data analysis.

**Availability and Implementation:** The ChIP-BIT2 package is available at http://sourceforge.net/projects/chipbitc/.

## Introduction

In order to identify transcription factor (TF) binding events or enrichment of histone markers (HMs), biologists run a sample ChIP-seq experiment to capture binding signals and background signals, and a second input ChIP-seq experiment to capture background signals only. The regulation strength of a TF or HM is often cell-type and condition specific. In some cases, the strength of a functional binding may be weak with a low read count observation. It is a challenging task to identify those weak binding events (with relatively low enrichment in the sample experiment but still much higher than that of input experiment) since they are more easily to be mixed with background signals also measured in the sample experiment. Recent studies have shown that functional effects of weak bindings can be very significant on gene transcription (1).

A number of ChIP-seq peak detection tools were developed using both sample and input ChIP-seq data (2–7). Read count is the most widely used signal format, which can be modeled to follow a Poisson distribution in sequencing data analysis (8–10). Although mathematically convenient to use read count, it gives rise to the difficulty in detecting weak binding events in ChIP-seq data because a ChIP-seq profile includes both binding and background signals and the medium or low read counts of weak binding events are often close to the level of background signals. Those tools developed based on read count are easily to classify weak binding events as background signals. Therefore, a Bayesian method, namely ChIP-BIT, has been proposed to reliably identify weak binding events at gene promoter regions (7).

Application of ChIP-BIT is limited since it can only be used to detect binding events of TFs at promoter regions. We present a new software package, ChIP-BIT2, to detect binding events of both strong and week signals from ChIP-seq data. ChIP-BIT2 expands the capability of ChIP-BIT for detecting binding events of TFs or HMs at any regions including gene promoter, enhancer or simply the whole genome. Moreover, ChIP-BIT2 is a C/C++ implementation. Compared to the original ChIP-BIT package (implemented in MATLAB), a significant speed improvement (~40%) can be gained for a pair of matched sample and input ChIP-seq profiles.

## Description

The challenge in detecting weak binding events lies in the ambiguity in differentiation of the binding signals from background signals. Background signal is not random noise. Its amplitude can be as high as a true binding signal. Using read intensity (log transform of average read coverage (7)), as illustrated in Fig. 1, ChIP-BIT2 will shrink the ‘distance’ between strong and weak binding signals and enlarge the ‘distance’ between weak and background signals, making it easier to detect weak bindings. The core functions of the algorithm are implemented in C/C++ as depicted in Fig. 2. ChIP-BIT2 can detect binding events either proximal to transcription starting sites (TSSs) or located at distal enhancer regions. As shown in Fig. 1 and Fig. 2, candidate ChIP-seq signal enriched regions are first identified through a data pre-processing step (with a similar strategy in PeakSeq (3)). Input background signal is then properly normalized against sample ChIP-seq signal. After loading annotated enhancer or promoter regions, ChIP-BIT2 can run in either ‘promoter mode’ or ‘enhancer mode’. Each promoter or enhancer region is first extended to a user-defined range and then partitioned into 200 bps bins. If no annotation region is provided, ChIP-BIT2 will detect all possibly enriched regions and partition the into bins as candidate regions. For each bin, sample read intensity, input read intensity and binding location (for promoter only) are calculated for each bin. A probability for binding occurrence is estimated using the Expectation-Maximization (EM) method. Consecutive bins with probabilities higher than a predefined threshold will be merged and finally outputted as enriched peaks. Using bin-based model, there is no need to set peak size limit, which makes ChIP-BIT2 flexible to detect narrow or wide peaks for TFs or HMs.

**Fig. 1.**
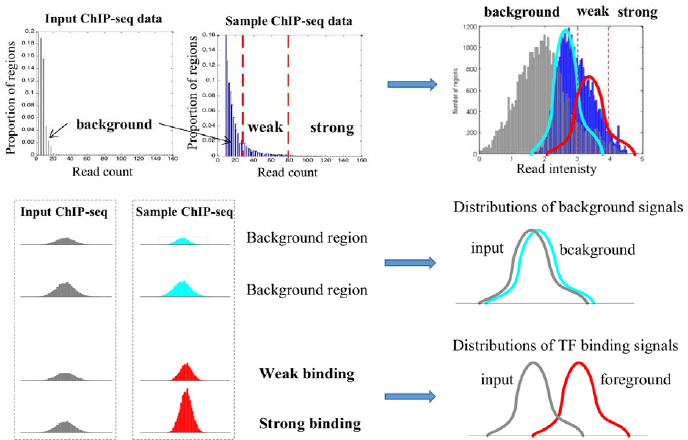
Converting read counts to read intensity for weak binding event detection.

**Fig. 2.**
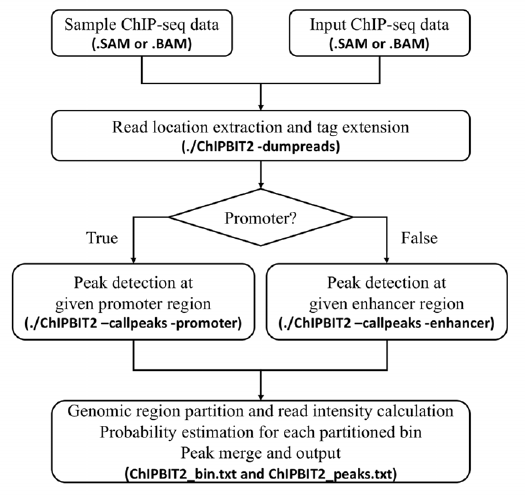
Workflow of ChIP-BIT2:

**Fig. 3.**
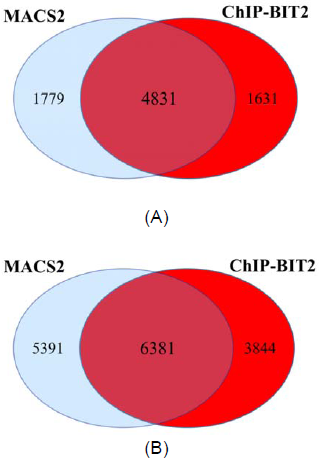
Venn diagrams of binding events detected by ChIP-BIT2 (red) and MACS2 (blue at 489 gene promoter regions (A) and 1050 enhancer regions (B).

## Results

We have compared ChIP-BIT2 and MACS2 using a breast cancer MCF-7 ChIP-seq data set with 39 TFs (as listed in Table 1). BAM files of selected TFs and their matched input data are downloaded from the ENCODE website (https://www.encodeproject.org/) and GEO database (https://www.ncbi.nlm.nih.gov/geo/). We used ChIP-BIT2 to process each TF and its match input data and predicted binding events at enhancer or gene promoter regions. Gene promoter regions were extracted from human reference genome hg19 as ±10k bps around each TSS. In total, we obtained 25,802 promoter regions regardless of potential overlap. Breast cancer MCF7 enhancer like regions were downloaded from ENCODE (https://www.encodeproject.org/data/annotations/). In total, there are non-overlap 33,957 enhancer regions (referred to hg19). We extended or pruned enhancer regions as ±1k bps around the original middle points. Finally, we used ChIP-BIT2 to call TFBSs at promoter and enhancer regions, respectively. ChIP-BIT2 and its previous MATLAB version were tested under CentOS Linux 7.3 on DELL T7600 workstation with 3.1GHz CPU (32 cores) and 128GB RAM. ChIP-BIT2 only uses one CPU core for one TF‘s ChIP-seq data processing and requires less than 2GB memory, so the running speed should be similar on any desktop or laptop with a similar CPU speed. The running speed of ChIP-BIT2 is summarized in Table. 2. Compared to its previous MATLAB version (supporting ‘promoter mode’ only), ChIP-BIT2 has gained a speed improvement of ~40%. Its speed in ‘enhancer mode’ is very similar to that in ‘promoter mode’, mainly determined by the number of reads in sample and input experiments.

We then compared ChIP-BIT2-detected peaks against those predicted by MACS2 (2) at selected promoter or enhancer regions associated with active genes (highly expressed) in MCF-7 cells. As shown in Fig. S3, a high proportion (73% for promoter or 54% for enhancer) of MACS2-detected binding events were captured by ChIP-BIT2. ChIP-BIT2 detected additional weak binding events (probability >0.9) together with those strong ones. Weak bindings at enhancer regions may play a more important role in gene regulation because an enhancer region is likely to be activated by a set of TFs, including both major activators and co-factors; the binding strength of cofactors may be weak (11). Therefore, ChIP-BIT2, by modeling read intensity instead of read count, can better eliminate background signals to effectively reduce false positive predictions, achieving an improved accuracy in detecting weak binding events.

## Conclusion

We have developed a software package, ChIP-BIT2, for detecting binding events of both strong and weak signals in promoter regions or enhancer regions. ChIP-BIT2, built upon a novel joint probabilistic model and implemented in C/C++, has been applied to a large set of ChIP-seq data and demonstrated its broad applicability in ChIP-seq data analysis for effective and reliable detection of binding events.

**Table S1.** 39 TF ChIP-seq profiles of breast cancer MCF-7 cells.

| Data source | TF symbols |
| --- | --- |
| ENCODE | CEBPB, CTCF, E2F1, EGR1, ELF1, EP300, FOSL2, FOXM1, GABPA, GATA3, HDAC2, JUND, MAX, MYC, NR2F2, NRSF, PML, POLR2A, RAD21, SIN3AK20, SRF, TAF1, TCF7, TCF12, TEAD4, ZNF217 |
| GSE26831 | c-FOS, c-JUN, FOXA1 |
| GSE41561 | CREB1, ER-α, KLF4, RXRA, TLE3 |
| GSE38901 | HSF1 |
| GSE44737 | MBD3 |
| GSE28008 | PBX1 |
| GSE22612 | TDRD3 |
| In house | NOTCH3 |

**Table S2.** Running speed of ChIP-BIT2 on peak detection for individual TFs

| TF symbols | ChIP-BIT2 Enhancer | ChIP-BIT2 Promoter | ChIP-BIT Promoter |
| --- | --- | --- | --- |
| KLF4 | 4m54 | 4m43 | 10m23 |
| TCF12 | 7m43 | 7m40 | 12m20 |
| c-JUN | 6m56 | 7m2 | 10m4 |
| c-FOS | 6m41 | 6m38 | 9m52 |
| JUND | 8m41 | 8m27 | 13m54 |
| RXRA | 8m46 | 8m37 | 13m10 |
| E2F1 | 9m59 | 9m48 | 16m53 |
| FOXA1 | 9m18 | 9m4 | 13m3 |
| RAD21 | 10m44 | 10m35 | 15m45 |
| TCF4 | 10m13 | 9m30 | 14m11 |
| TLE3 | 13m46 | 11m59 | 22m33 |
| ZNF217 | 10m36 | 10m29 | 15m11 |
| HSF1 | 11m17 | 6m25 | 14m55 |
| ELF1 | 12m37 | 12m29 | 17m42 |
| ER-α | 13m41 | 12m29 | 20m10 |
| POL2A | 12m29 | 11m34 | 16m47 |
| NOTCH3 | 14m25 | 13m38 | 21m7 |
| SRF | 15m30 | 13m35 | 22m50 |
| TDRD3 | 25m20 | 23m21 | 48m44 |
| CEBPB | 13m50 | 13m42 | 21m31 |
| FOSL2 | 14m18 | 13m8 | 18m50 |
| GATA3 | 14m3 | 14m5 | 20m56 |
| GABPA | 13m41 | 13m29 | 18m18 |
| NR2F2 | 15m15 | 14m31 | 21m38 |
| PBX1 | 17m11 | 15m21 | 25m47 |
| CTCF | 14m35 | 13m59 | 19m24 |
| EP300 | 14m43 | 14m15 | 18m38 |
| FOXM1 | 15m11 | 14m25 | 19m25 |
| PML | 14m34 | 9m26 | 19m40 |
| CREB1 | 17m19 | 16m43 | 24m43 |
| HDAC2 | 14m34 | 15m36 | 21m34 |
| TEAD4 | 14m50 | 14m11 | 19m35 |
| EGR1 | 16m32 | 15m44 | 20m39 |
| NRSF | 17m13 | 16m27 | 22m12 |
| SIN3A | 19m29 | 15m0 | 27m15 |
| MYC | 31m32 | 28m7 | 46m9 |
| MAX | 20m44 | 19m55 | 26m35 |
| TAF1 | 20m7 | 19m33 | 25m50 |
| MBD3 | 27m39 | 26m47 | 36m6 |

